# Invertebrate species produce taxon-specific acoustic profiles under controlled conditions

**DOI:** 10.64898/2026.04.12.718076

**Authors:** Amy Annells, Martin F. Breed, Timothy R. Cavagnaro, Riley J. Hodgson, Sofie Costin, Tarryn Davies, Alex Taylor, Jake M. Robinson

## Abstract

Soil degradation threatens food security, climate regulation, biodiversity and human wellbeing worldwide. Up to 75% of the world’s soils are already degraded, and in response, global restoration efforts are rapidly scaling up to meet international targets. Monitoring soil biodiversity recovery remains a major barrier to tracking restoration success, particularly for invertebrates that underpin key processes including nutrient cycling and soil aggregation. Traditional sampling methods are labour-intensive, destructive and poorly suited to long-term or landscape-scale monitoring. Soil ecoacoustics is rapidly emerging as a promising non-destructive soil biodiversity monitoring tool. However, its capacity for taxonomic resolution of invertebrate groups remains untested. Here, we present a proof-of-concept study that establishes the potential for a soil invertebrate acoustic classifier. We used a low-cost, sound-attenuated recording system and quantified 19 spectral and temporal audio features from six morphologically and behaviourally distinct invertebrate species under controlled conditions. Acoustic profiles differed among taxa and generally clustered into podous and apodous groups (i.e., organisms with legs versus those without). Variation was driven primarily by taxon identity rather than body mass, suggesting that acoustic signatures capture taxon-specific traits. This work provides a foundation for developing automated acoustic classifiers that could enable scalable, non-destructive soil biodiversity monitoring.

## 1. INTRODUCTION

Healthy soils are multifunctional living systems. They are foundational to terrestrial life, supporting 98% of calories consumed by humans [1] and are home to 59% to >99.9% of Earth’s species [2, 3]. Currently, 75% of the world’s soils are affected by degradation, a figure that could rise to 90% by 2050 if harmful practices persist [4]. In response to this growing global ecological crisis, the United Nations declared 2021-2030 the Decade on Ecosystem Restoration [5], spurring a call to action and urgency in restoring degraded ecosystems. Healthy soils support many ecological functions and processes, including plant productivity, water regulation, and a vast belowground biodiversity that sustains life aboveground [6]. Among this belowground diversity, soil fauna forms a critical component of restoration success. Indeed, invertebrates play an important role in overall soil health, as they contribute to nutrient cycling, soil aggregation, and can influence plant health and food web dynamics [7].

Effective biodiversity monitoring underpins successful restoration by enabling the assessment of intervention outcomes, the early detection of persistent or emerging disturbances and the iterative refinement of actions through adaptive management [8]. However, detecting, identifying and monitoring soil biota poses numerous challenges. Traditional methods (e.g., pitfall traps, soil sampling, DNA barcoding, and Berlese funnel approaches) often have negative ecological impacts and are laborious and costly at large scale [9, 10]. Therefore, developing informative, low-cost and accessible tools that overcome prior limitations to monitor soil health is imperative if we want to develop efficient, rapidly deployable, and scalable tools [11]. This need for innovation has led to new developments, including acoustic monitoring technologies [12].

For decades, passive acoustic monitoring (PAM) has been widely applied in bioacoustics research across a range of vertebrate taxa, such as bats, birds, frogs and marine mammals, with relatively high efficacy [13]. More recently, ‘ecoacoustics’ (the investigation of natural and anthropogenic sounds and their relationships with the environment) has been developed for soils [14, 15]. Soil soundscape indices (e.g., acoustic complexity index, bioacoustic index, acoustic diversity index) can reflect progress towards restoration targets under field conditions [16, 17]. However, soil ecoacoustic indices provide only a coarse overview of biodiversity, and it remains unclear whether acoustic approaches can resolve specific invertebrate taxa from their signals [18].

Although acoustic monitoring has been applied to invertebrates, its application remains limited and highly species-specific [19]. Most studies focus on detecting the presence or absence of pest invertebrates (e.g., red palm weevil *Rhynchophorus ferrugineus*, larvae of Scarabaeidae [20, 21]) that provide limited insight into the broader community complexity that would emerge from simultaneously investigating multiple invertebrate taxa. Where multiple taxa have been studied, efforts have largely focused on soniferous or flying invertebrates (e.g., field crickets *Gryllus bimaculatus*, cicadas (Cicadidae) or bees (Apoidea) [22, 23]), whose acoustic signals (e.g., wing buzzing, stridulation, or chirping) are captured by existing above-ground recording technologies. Conversely, the low-frequency, substrate- or leaf litter-borne vibrations and subtle soil-transmitted sounds produced by belowground and near-surface dwelling taxa (e.g., worms (Annelida), ants (Formicidae), and spiders (Araneae)) remain poorly characterised.

Some studies have attempted to use publicly available invertebrate sound libraries; however, these datasets often introduce confounding variability due to inconsistent recording durations, environmental conditions and equipment types [24]. As such, there is a growing need for a standardised acoustic classifier tailored to soil ecosystems that can detect and differentiate between soil-dwelling invertebrate taxa. Such a tool could enhance our ability to interpret belowground and surface-dwelling biodiversity patterns, monitor restoration progress and advance the emerging field of soil ecoacoustics.

Here, we take a critical early step toward developing an acoustic classifier for soil invertebrates. Our goal was to undertake a proof-of-concept study that tests whether common soil and litter-dwelling invertebrates produce taxon-specific acoustic profiles under controlled conditions. We used a low-cost, standardised recording system and quantified spectral and temporal audio features from six readily available invertebrate species that are morphologically and behaviourally distinct (i.e., house cricket *Acheta domesticus*, garden snail *Cornu aspersum*, red wiggler worm *Eisenia fetida*, lobster cockroach *Nauphoeta cinerea*, huntsman spider *Pediana horni*, and mealworm *Zophobas morio (*larval stage of a aarkling beetle). We hypothesised that: (i) taxa would exhibit distinct acoustic profiles, (ii) acoustic differentiation would reflect morphological traits (e.g., podous or apodous), and (iii) taxa identity rather than body mass would explain most acoustic variation.

## 2. METHODS

### (a) Experimental setup

To create a standardised environment for recording invertebrate movement and acoustic signals, we used a custom-built sound-attenuated chamber (Figure 1*a*). For construction, the interior of a 54 L plastic container was lined with sound attenuation foam to minimise ambient noise. A lipped aluminium plate (30 × 45 cm; 1 mm thickness) was positioned within the chamber to serve as a standardised surface for invertebrate detection. For recording invertebrate movement, we utilised a cost-effective recording assembly (<AUD$1500), enabling our methodologies to be readily applied in other ecological systems. A JrF C-Series Pro contact microphone sensor (manufacturer: www.jezrileyfrench.co.uk; Yorkshire, U.K.) was affixed beneath the centre of the aluminium plate using silver adhesive tape. The C-series contact microphones are more sensitive to audio vibrations through contact with solid objects, making them well-suited for detecting the movement of invertebrates across the aluminium plate. In addition, they offer a wide frequency response, particularly optimised for the low-end and mid-frequency ranges [17, 25]. The contact microphone was connected to a micBooster pre-amp and connected to an acoustic recording device (Zoom F6 multitrack field recorder; Designed by ZOOM, Tokyo, Japan). To prevent invertebrates from escaping the chamber, an acrylic sheet (31 × 46 cm; 3 mm thickness) was placed over the aluminium plate. Concurrent video footage was recorded via a Samsung Galaxy S25+ camera mounted above the sound attenuation chamber on a microphone stand, which enabled visual verification of invertebrate movements for correlation with acoustic activity.

**Figure 1.**
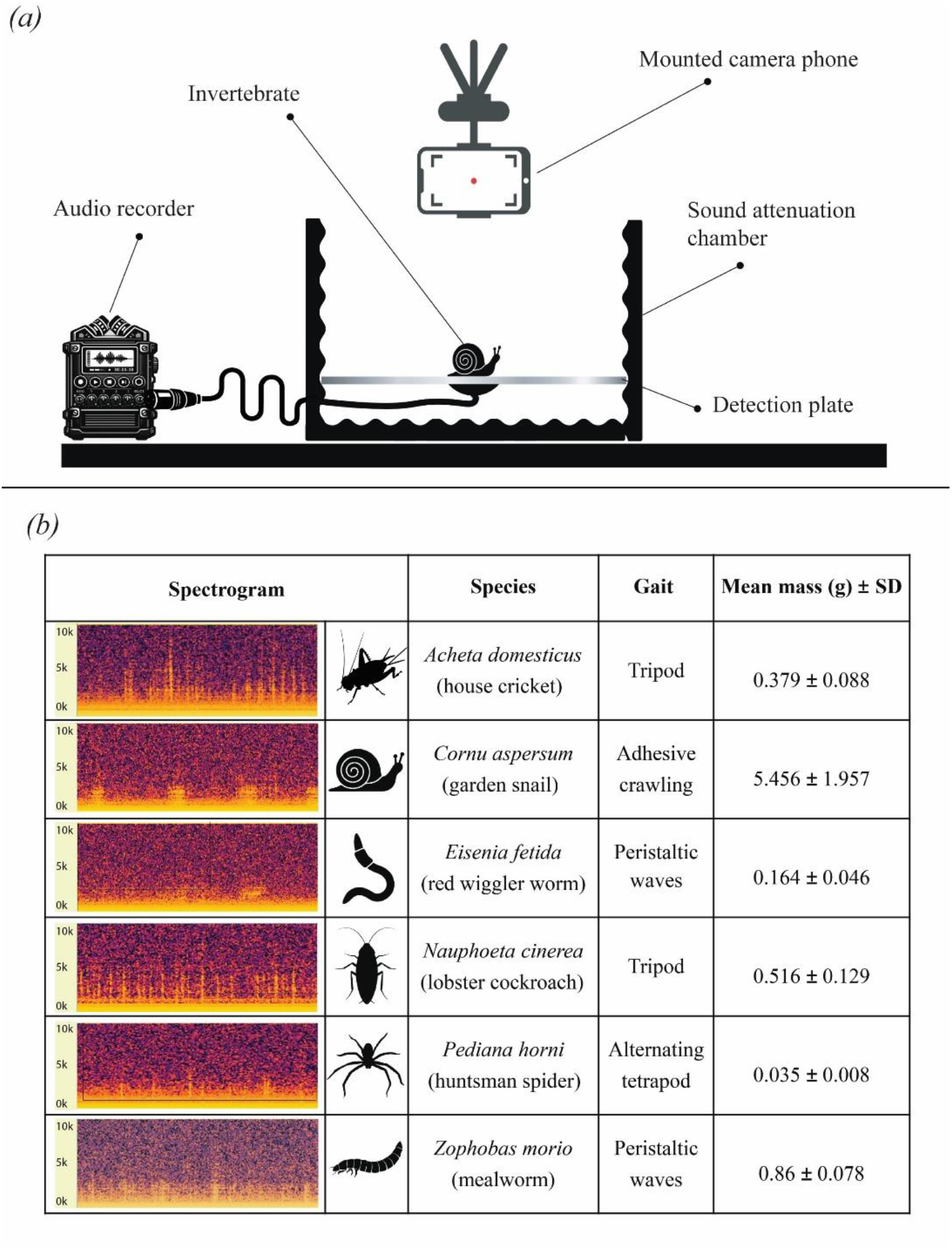
**a)** recording of soil invertebrates was achieved using a low-cost custom-built sound chamber where a 50 cm plastic container lined with sound attenuation foam contained a 1 mm lipped aluminium plate with a contact microphone connected to the acoustic recording device and an overhead phone camera. **b)** Overview of the 6 invertebrate species studied, spectrogram images sourced from Audacity® (version 3.7.7) files of invertebrate recordings, frequency in Hz [26].

Six locally common and readily accessible invertebrate species were chosen to capture differences in gait, morphology, and taxonomic class (Figure 1*b*). For each species, 10 individuals were randomly selected for acoustic analysis from larger population’s (<50 individuals). However, *P. horni* was constrained to 10 individuals total due to supplier limitations. The body mass of each invertebrate was measured using an analytical scale immediately prior to recording. During each recording, the invertebrate was placed at the centre of the aluminium plate and allowed to move freely for five minutes while acoustic data were collected. To account for ambient background noise, a five-minute control recording was obtained prior to each of the recording trials for each species. We recorded Waveform Audio Format (.WAV) sound files at a 24-bit depth and a sampling rate of 48 kHz on the Zoom F6 units. We used +60 dB preamp gain based on pilot tests to maximise the signal-to-noise ratio (SNR) for quiet, substrate-borne signals while avoiding clipping (24-bit, peaks −12 to −3 dBFS; 0% clipped samples). Audio recordings were saved as.WAV files and used for downstream analysis. All recordings were conducted over three sessions in a temperature and sound-controlled recording booth at Flinders University, South Australia.

### (b) Audio data pre-processing and feature extraction

To extract invertebrate audio profiles, we implemented a feature extraction pipeline in Python (v 3.13.2) using the following libraries: ‘os’, ‘numpy’ [27], ‘pandas’ [28], ‘librosa’, and ‘scipy.signal’[29]. This pipeline was used to extract 19 key audio features (Supplementary Table 1). All recordings (.WAV format) were converted to mono to standardise downstream analysis by generating a single signal for consistent computation, and at a sampling rate of 48 kHz to capture the biological frequency ranges of invertebrates. The audio signals were then amplitude-normalised and trimmed by 10s at the start and the end to remove handling artifacts before analysis. We then used the ‘librosa’ library to extract spectral features of the normalised audio files [30]. Thirteen Mel-frequency cepstral coefficients (MFCCs; features that summarise the spectral shape of a sound based on human-like frequency perception) were computed from each signal and averaged across frames. The use of 13 MFCCs is common in audio signal processing as it balances information and computational efficiency; the first 12 coefficients capture the shape of the spectral envelope, and the 13^th^ is the overall slope [31]. We used ‘librosa’ library to extract additional spatial and temporal audio descriptors from each audio file, including average inter-pulse interval, spectral centroid, spectral bandwidth, zero-crossing rate, and root-mean-square energy. To reduce bias and capture peak frequency patterns from the audio files, a 2-12 kHz band-pass filter was applied, reducing low-frequency background noise (e.g., air conditioning hum). The filtered waveform was then short-time Fourier transformed (STFT) to generate magnitude spectrograms for temporal (∼12 ms) and spatial (21.5 Hz/bin) balance for short pulsating sounds generated by invertebrate movement. Frames 35 dB below the median energy were filtered out to further reduce background noise and conflation from silent frames, with the dominant frequency then calculated from the retained frames. The extracted features were compiled into a single dataset and exported as a comma-separated file (.) for statistics.

### (c) Statistical analysis

All downstream statistics were done in R (version 4.5.3) using the R Studio interface (v 4.5.1) [32]. For data visualisations, a combination of R and Adobe Illustrator Creative Cloud 2025 (v 6.8.0.821) [33] was used.

### Invertebrate audio profiles

To compare invertebrate audio profiles we used Principal Component Analysis (PCA) on the filtered, scaled dataset via the *prcomp* function in base R. First, highly correlated (r>0.70) acoustic parameters were identified and removed using *findCorrelation* (Spearman) in caret [34] to reduce multicollinearity among variables, and the remaining variables were scaled (z-transformed). Following PCA construction [35], we used 95% confidence interval ellipses to represent within-species variation. Acoustic variable loadings were then extracted from the PCA matrix to identify the features contributing most strongly to the ordination axes, with the top six variables plotted as biplot arrows on the PCA ordination. To investigate whether the overall acoustic profile differed significantly amongst each species, we used a permutational multivariate analysis of variance (PERMANOVA) (9,999 permutations) via *adonis2* in *vegan* [36]. Pairwise PERMANOVA were then conducted for significant tests using the *pairwise*.*adonis2* function in *vegan* [36] with p-values adjusted for multiple comparisons using the Benjamini–Hochberg procedure to control the false discovery rate [37]. Homogeneity of multivariate dispersion among species was also assessed using permutational analysis of multivariate dispersions (PERMDISP) [38] on a Euclidean distance matrix derived from the PCA scores. Pairwise comparisons were again, adjusted with the Benjamini-Hochberg FDR correction [37].

To address non-normality and heteroscedasticity in the relationships between invertebrate species and each of the 19 audio parameters, we used randomised one-way ANOVAs with 10,000 permutations. Randomised (permuted) two-sample t-tests were then applied to estimate pairwise differences 10,000 times, among species for each audio parameter, and multiple comparisons used the Benjamini-Hochberg FDR correction [37].

### Invertebrate mass

Redundancy analysis was used to determine whether mass influenced the invertebrate audio profile, once species effects were accounted for. Prior to ordination, highly correlated (>0.70) acoustic parameters were identified and removed using the *findCorrelation* function in caret [34] to reduce multicollinearity among predictors, and then the remaining variables were scaled (z-transformed). Using species and body mass as explanatory variables, a global redundancy analysis was then fitted using the *rda* function in *vegan* [36]. The significance of the model was determined using permutational ANOVAs (10,000 permutations) on both the overall model and the marginal terms to determine the contribution of each explanatory variable. Redundancy analysis site (sample) scores for the first two canonical axes were extracted visualised, with 95% confidence interval ellipses [35].

## 3. RESULTS

The acoustic profiles of the six invertebrate species differed strongly (PERMANOVA: *df* = 5, *f* = 11.031, *R*^*2*^ = 0.505, *p* < 0.001; Fig. 2). Differences in multivariate dispersion were detected (*betadisper p* = 0.007), driven by cockroaches (Supplementary Figure 1). Pairwise permutational comparisons confirmed that nearly all species differed from one another (*p* < 0.05) and from control recordings (*p* < 0.05), except garden worms and snails (*p* = 0.128). Distinct clustering was evident between podous (legged) and apodous (legless) taxa, with cockroaches and spiders separated along PC1. Crickets showed partial overlap across multiple taxa, while the apodous species (i.e., garden worms, snails, and mealworms) clustered closely with one another, suggesting that invertebrates lacking appendages may produce more similar acoustic profiles. Five of the six most influential acoustic parameters (i.e., MFCC 3, MFCC 5, MFCC 9, MFCC 12, and MFCC 13) were strongly aligned with the first axis, indicating that these spectral features were major drivers of interspecific acoustic differentiation.

**Figure 2.**
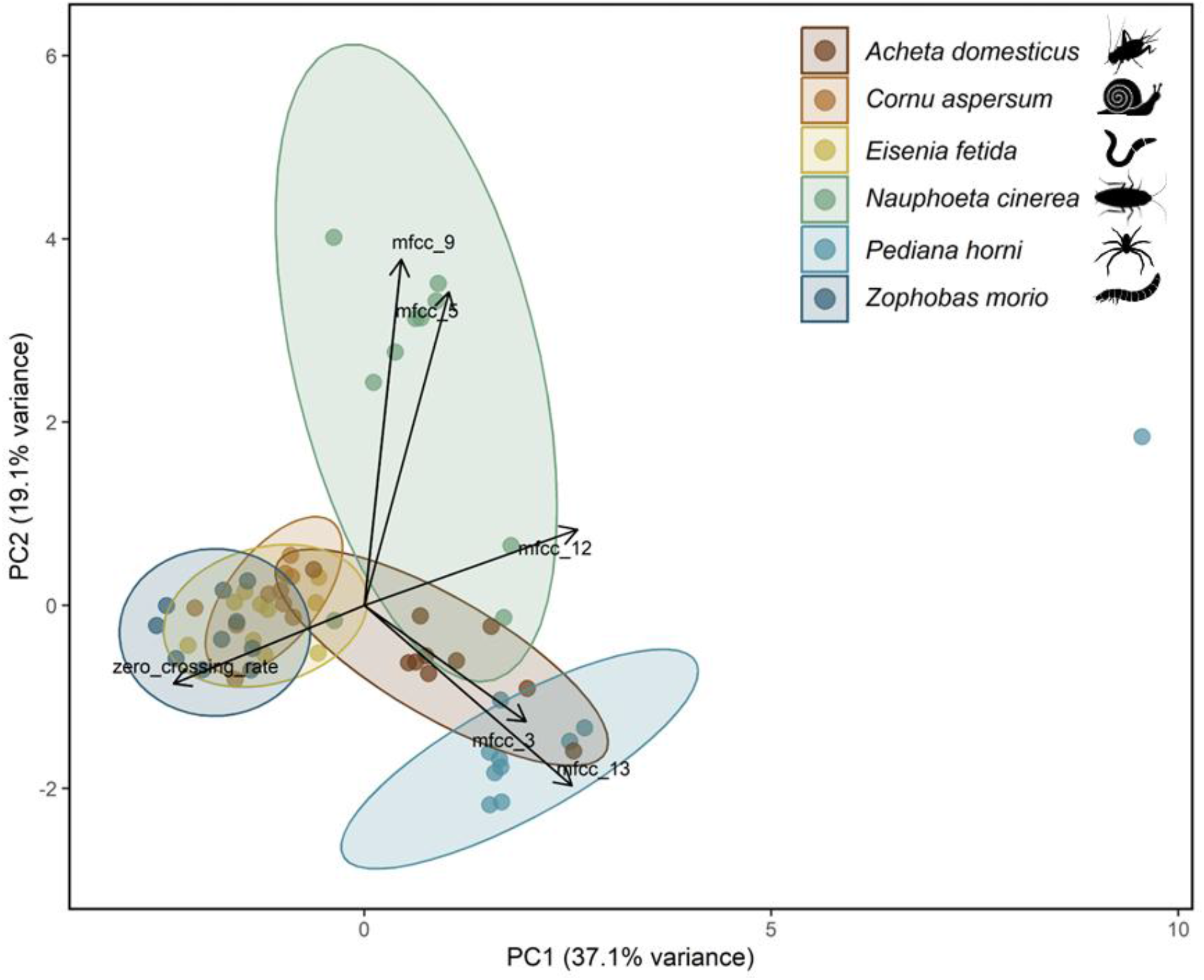
Soil invertebrates produce distinct acoustic profiles. Principal component analysis (PCA) of uncorrelated, z-transformed audio features, with 95% confidence ellipses. Each point represents an individual, with proximity indicating similarity in acoustic profiles. *PERMANOVA* \*\**p* = 0.001.

Species differed in 18 of the 19 acoustic features (randomised one-way ANOVA, 10,000 permutations; *p* < 0.001 for each), where RMS energy did not vary by species (*p* = 0.627) (Figure 3). For several parameters, effects were restricted to contrasts between podous and apodous taxa, consistent with the separation observed in the PCA ordination (Fig. 2). Individual animal mass had no effect on invertebrate audio profiles after accounting for species identity (RDA *f* = 0.309, *p* = 0.878; Supplementary Figure 2).

**Figure 3.**
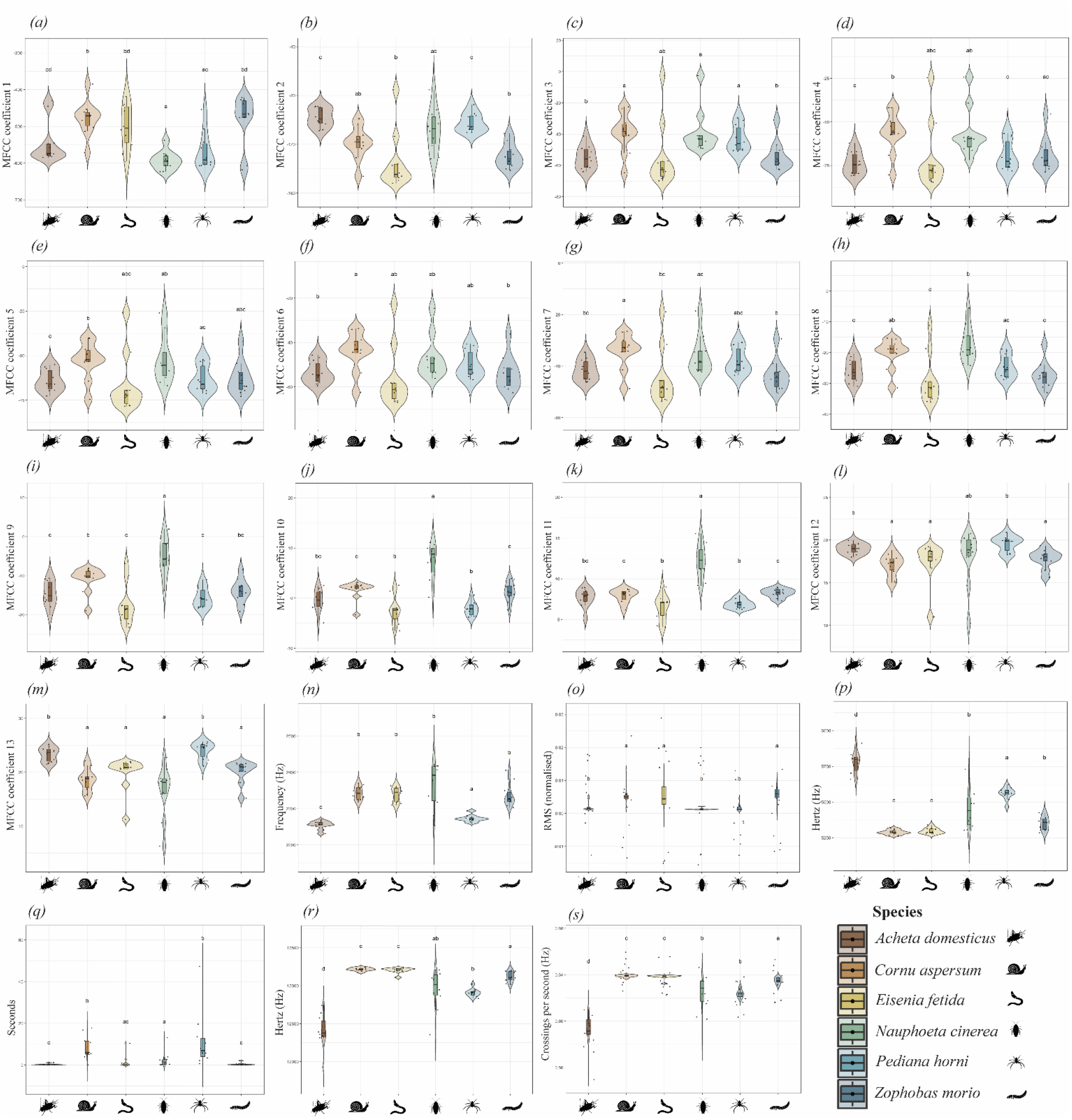
Specific audio parameters may be sensitive to taxa-specific morphology (y-axis is scaled independently). a–m: MFCC 1–13 (Mel-frequency cepstral coefficients; descriptors of spectral shape), n: Peak frequency (frequency with highest energy), o: RMS energy (overall signal amplitude), p: Spectral bandwidth (spread of frequencies around the centre), q: Average interpulse interval (time between successive pulses), r: Spectral centroid (centre of mass of the spectrum), s: Zero-crossing rate (rate of signal sign changes). Letters within each plot denote significant differences based on post-hoc comparisons (groups sharing letters are not significantly different).

## 4. DISCUSSION

Here we show that six common soil- and litter-dwelling invertebrates produced distinct and quantifiable acoustic profiles under controlled conditions, supporting our first hypothesis. Using a low-cost recording system and a reproducible feature-extraction pipeline, we demonstrate that multivariate combinations of spectral and temporal features discriminate taxa with contrasting morphologies and locomotor strategies. Species clustered strongly, with clear separation between podous and apodous taxa, supporting our second hypothesis. Once species effects were accounted for, body mass had little explanatory power, providing limited evidence for our third hypothesis. These results indicate that morphological and behavioural traits drive acoustic differentiation, not simple body mass scaling relationships. These findings provide empirical evidence that soil invertebrate taxa exhibit taxa-specific acoustic signatures detectable in a controlled environment. By demonstrating that species identity can be statistically resolved from substrate-borne signals, this work advances soil ecoacoustics beyond aggregate soundscape indices toward taxonomic resolution. In doing so, it lays the empirical foundation for automated acoustic classifiers capable of supporting scalable, non-destructive monitoring of soil biodiversity in restoration contexts.89

We show strong interspecific clustering of acoustic signals, highlighting that acoustic-based identification of soil-associated invertebrates is plausible. While feature-rich, classifier-based approaches are emerging in soil ecoacoustics, studies still rely heavily on summary indices that aggregate soundscape properties without taxonomic resolution [39]. Existing approaches, such as the Acoustic Complexity Index (ACI) and Acoustic Diversity Index (ADI), have successfully captured community-level acoustic activity and linked it to soil biological richness [16, 17, 40], but they cannot attribute signals to particular taxa or processes. Our study aligns with current opinions of soundscape level indices, such as that of Metcalf *et al*. (2024) [39], who described the need to undertake ex-situ mesocosm trials of known soil faunal communities to better understand the identifying characteristics of species’ acoustic signals. The interspecific clustering observed in our study demonstrates that species-level discrimination is achievable under controlled conditions and lays the groundwork to expand acoustic identification of soil invertebrates to species or functional group resolution. Such a resolution would have broad implications for how biodiversity and ecosystem function are monitored. In restoration contexts, soil invertebrate monitoring is often limited (e.g., destructive sampling methods) [38] or there is difficulty in determining source viability in eDNA technologies (i.e. alive, dead or transient) [41]. As acoustic profiles must come from an active invertebrate it could overcome these disadvantages and, be applied to track colonisation trajectories of functionally important taxa (e.g., decomposers, detritivores) and detect shifts in community structure following changes in vegetation cover or microclimate. Such advantages would not only have implications for soil invertebrate monitoring in restoration contexts but could be expanded to all guises of invertebrate monitoring (i.e. agricultural pest management and biosecurity, species conservation).

Distinct clustering of podous and apodous invertebrates within ordination space supports our hypothesis that morphological traits strongly influence acoustic signatures. Consistent with this pattern, body mass had little explanatory power once species identity was accounted for, reinforcing that trait differences rather than simple size scaling underpin acoustic divergence. These results are consistent with previous invertebrate acoustic studies. For example, Rodriguez et al. [39] showed that bee wing morphology influences acoustic signatures, with longer wings associated with lower frequencies, while Escola et al. [40] demonstrated that species-specific behavioural patterns and abdominal movements in cicadas enable distinct acoustic identification.

We observed that podous taxa with broader behavioural repertoires, particularly cockroaches and, to a lesser extent, crickets, drove multivariate acoustic dispersion. These taxa often produce complex mechanical signals, including stridulation, leg tapping, and general locomotor activity [42], which may contribute to higher within-species variation. In contrast, apodous taxa, such as snails, exhibit a simpler acoustic repertoire, generating vibrations primarily through adhesive locomotion, which may result in lower within-species variation [43]. Although 18 of 19 audio parameters differed among species, the discriminatory power of individual features was context dependent. Mels frequency cepstral coefficients, which capture the short-term spectral envelope, showed broad interspecific differentiation and contributed strongly to ordination structure. In contrast, spectral bandwidth did not show clear morphology-based clustering, suggesting that some audio parameters are more sensitive to taxa-specific morphology than others. These results underscore the necessity of a multivariate classification framework, as no single parameter adequately characterises soil invertebrates. Integrating cepstral, spectral, temporal and energy metrics will be critical for machine learning applications, where combined MFCC and complementary feature sets have been shown to improve automated insect detection accuracy relative to single-feature approaches [44].

While encouraging, our study represents an early proof-of-concept, and several limitations should guide future work. First, recordings were obtained in a controlled environment on a homogeneous aluminium substrate with minimal background noise. Litter and soils differ markedly in structure, compaction and moisture, all of which influence sound propagation [12]. Future studies should progressively increase environmental complexity, for instance, adding soil or leaf litter layers and then advancing to controlled mesocosm experiments. Second, individuals were recorded in isolation to obtain discrete acoustic profiles. Soil communities contain many interacting taxa occupying diverse niches [45], and signal discrimination will therefore be more complex under field conditions. Experiments using mock communities may be required to test classification performance in ecologically realistic assemblages before testing under field conditions against more established soil biodiversity assays (i.e., pitfall surveys, COI amplicons etc.). Despite these constraints, our results provide foundational evidence that soil invertebrate assemblages can be differentiated acoustically. This establishes a necessary first step toward automated classifiers capable of resolving fine-scale biodiversity patterns in soil ecosystems.

## 5. Conclusion

This study provides empirical evidence under controlled conditions that common soil and litter-dwelling invertebrates produce distinct and statistically discriminable acoustic profiles. Taxa identity, rather than body mass, explained most acoustic variation, indicating that morphological and behavioural traits underpin acoustic differentiation. By showing that taxa-specific signatures can be resolved from substrate-borne signals, our study helps move soil ecoacoustics beyond aggregate soundscape indices toward taxonomic resolution. This establishes a clear foundation to develop automated, soil-specific acoustic classifiers. Such tools would enable non-destructive, scalable and continuous soil fauna monitoring, strengthening our capacity to track biodiversity recovery and evaluate restoration outcomes at scale.

## ACKNOWLEDGEMENTS

We acknowledge the Kaurna People, the traditional owners of the lands of the Tarntanya (Adelaide) region in Australia, on which this study was undertaken.

