## Supplementary Table 1 for "Invertebrate species produce taxon-specific acoustic profiles under controlled conditions"

**SUPPLEMENTARY DATA
Supplementary Table S1:** Overview of audio parameters analysed

| ***Acoustic Parameter*** | ***Acronym*** | ***Description*** | ***Variables Used*** |
| --- | --- | --- | --- |
| Mel Frequency Cepstral Coefficients | MFCC | A compact representation of the spectral properties of a sound signal, specifically designed to mimic how humans perceive audio. They summarize the spectral envelope of short sound frames using a scale that reflects human sensitivity to different frequencies (the Mel scale). | **Frequency** **content** (Spectrum), **Amplitude** (via energy in frequency bins), **Time** (in form of short frames), **Perceptial** **scaling** (mel scale) |
| Spectral Centroid | SC | Indicates the "center of mass" of a sound spectrum. It reflects where the average frequency is located in the power spectrum — higher centroid values generally correspond to brighter, sharper, or more high-pitched sounds, while lower values indicate duller, bass-heavy sounds. | **Frequency** (location of each spectral bin), **Amplitude** (magnitude at each frequency bin), **Time** (applied frame-by-frame for temporal changes) |
| Spectral Bandwidth | SBW | Measures the spread of the frequency content around the spectral centroid. It quantifies how "wide" or "narrow" the frequency energy is distributed, offering insight into the complexity or noisiness of a sound. Narrow bandwidth indicates tonal, focused energy (like a whistle), while wide bandwidth reflects broadband signals (like wind or insect choruses). | **Frequency** (locations of spectral bins), **Amplitude** (used for weighting), **Time** (to observe changes across frames), **Spectral** **Centroid** (central reference point) |
| Zero Crossing Rate | ZCR | Measures how often the amplitude of an audio signal crosses the zero line — in other words, how frequently the waveform changes sign (from positive to negative or vice versa). It’s a useful indicator of signal noisiness or sharpness and is especially helpful for distinguishing between tonal and atonal or noisy sounds. | **Amplitude** (particularly the sign of amplitude values), **Time** (sample-by-sample transitions) |
| Root Mean Square Energy | RMS | Measures the average power or energy of an audio signal over time. It represents how strong or loud a signal is, smoothing out peaks and giving a more perceptually relevant value than raw amplitude. It is particularly useful in assessing signal intensity and dynamics, such as loud vocalizations, sudden events, or changes in ambient acoustic activity. | **Amplitude** (raw signal magnitude, squared and averaged), **Time** (frame-by-frame temporal tracking) |
| Average Pulse Interval | API | Measures the mean time gap between successive pulses (i.e., discrete bursts of sound energy) in a signal. It reflects the temporal rhythm or pacing of acoustic events and is especially useful in analyzing species that produce patterned or periodic sounds, such as cricket chirps or frog calls. | **Amplitude** (for detecting discrete pulses), **Time** (to measure gaps between pulses) |
| Peak Frequency | PFrq | Refers to the frequency component within a sound that carries the highest amplitude (or power) at a given moment or over a defined window. It is often used to identify the most prominent tonal feature of a signal and can help differentiate species or events based on their dominant pitch or resonant characteristics. | **Frequency** (location of each spectral bin), **Amplitude** (used to determine which frequency is dominant), **Time** (to track peak frequency across a recording) |

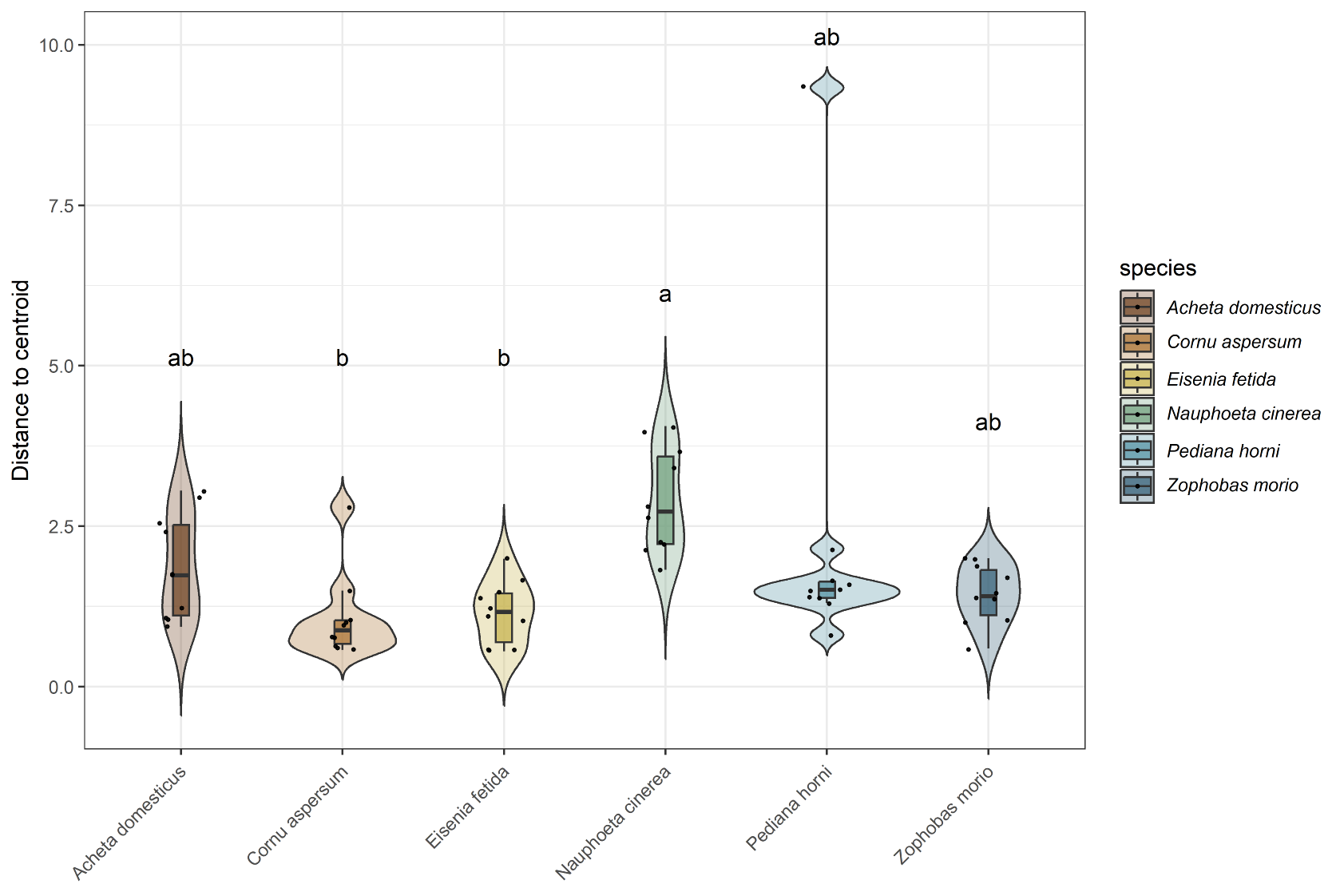

**Supplementary Figure S1:** Dispersion is driven by podous invertebrates**.** Distance to centroid showing homogeneity of multivariate dispersion among invertebrate taxa. Letters denote significant differences based on post-hoc comparisons (groups sharing letters are not significantly different).

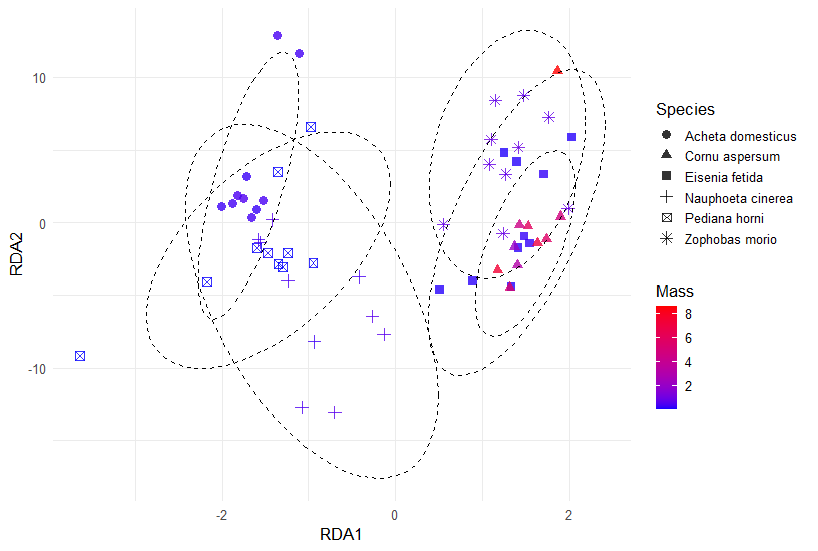

**Supplementary Figure S2 UPDATE :** Mass has little effect on audio profiles of soil associated invertebrates. Redundancy analysis on invertebrate acoustic profiles controlling for species. Colour-scaled by body mass and shaped by species.
